# Bacterial microcompartments linked to the flavin-based extracellular electron transfer drives anaerobic ethanolamine utilization in *Listeria monocytogenes*

**DOI:** 10.1101/2020.10.27.358424

**Authors:** Zhe Zeng, Sjef Boeren, Varaang Bhandula, Samuel H. Light, Eddy J. Smid, Richard A. Notebaart, Tjakko Abee

**Affiliations:** Food Microbiology, Wageningen University and Research, Wageningen, Netherlands; Laboratory of Biochemistry, Wageningen University and Research, Wageningen, Netherlands; Department of Molecular and Cell Biology, University of California, Berkeley, CA, USA; Department of Microbiology, The University of Chicago, Chicago, IL, USA

## Abstract

Ethanolamine (EA) is a valuable microbial carbon and nitrogen source derived from phospholipids present in cell membranes. EA catabolism is suggested to occur in so-called bacterial microcompartments (BMCs) and activation of EA utilization (*eut*) genes is linked to bacterial pathogenesis. Despite reports showing that activation of *eut* in *Listeria monocytogenes* is regulated by a vitamin B12-binding riboswitch and that upregulation of *eut* genes occurs in mice, it remains unknown whether EA catabolism is BMC dependent. Here, we provide evidence for BMC-dependent anaerobic EA utilization via metabolic analysis, proteomics and electron microscopy. First, we show B12-induced activation of the *eut* operon in *L. monocytogenes* coupled to uptake and utilization of EA thereby enabling growth. Next, we demonstrate BMC formation in conjunction to EA catabolism with the production of acetate and ethanol in a molar ratio of 2:1. Flux via the ATP generating acetate branch causes an apparent redox imbalance due to reduced regeneration of NAD+ in the ethanol branch resulting in a surplus of NADH. We hypothesize that the redox imbalance is compensated by linking *eut* BMC to anaerobic flavin-based extracellular electron transfer (EET). Using *L. monocytogenes* wild type, a BMC mutant and a EET mutant, we demonstrate an interaction between BMC and EET and provide evidence for a role of Fe^3+^ as an electron acceptor. Taken together, our results suggest an important role of anaerobic BMC-dependent EA catabolism in the physiology of *L. monocytogenes*, with a crucial role for the flavin-based EET system in redox balancing.

**IMPORTANCE:** *Listeria monocytogenes* is a food-borne pathogen causing severe illness and, as such, it is crucial to understand the molecular mechanisms contributing to pathogenicity. One carbon source that allows *L. monocytogenes* to grow in humans is ethanolamine (EA), which is derived from phospholipids present in eukaryotic cell membranes. It is hypothesized that EA utilization occurs in bacterial microcompartments (BMCs), self-assembling subcellular proteinaceous structures and analogs of eukaryotic organelles. Here, we demonstrate that BMC-driven utilization of EA in *L. monocytogenes* results in increased energy production essential for anaerobic growth. However, exploiting BMCs and the encapsulated metabolic pathways also requires balancing of oxidative and reductive pathways. We now provide evidence that *L. monocytogenes* copes with this by linking BMC activity to flavin-based extracellular electron transfer (EET) using iron as an electron acceptor. Our results shed new light on an important molecular mechanism that enables *L. monocytogenes* to grow using host-derived phospholipid degradation products.

## Introduction

Pathogens have evolved mechanisms to utilize specific metabolites as carbon-sources to sidestep nutritional competition with commensal bacteria in the human gastrointestinal tract [3-6]. Ethanolamine(EA), a product of the breakdown of phosphatidylethanolamine from eukaryotic cell membranes, is such a metabolite and is abundant in the human gastrointestinal tract [7, 8]. It has been shown that some species in the gastrointestinal tract like *Salmonella enterica, Enterococcus faecalis, Clostridium perfringens* can use EA as carbon source while for some other human pathogens including *Listeria monocytogenes*, the putative use of EA as a substrate was postulated based on the presence of a similar gene cluster [7-9]. The capability to utilize EA is encoded by the ethanolamine utilization (*eut*) operon [8-10]. EA is converted to acetaldehyde and ammonia by ethanolamine ammonia lyase EutBC [8, 11, 12]. Acetaldehyde can be catabolized to ethanol by the alcohol dehydrogenase EutG [10] or to acetyl-CoA, by acetaldehyde dehydrogenase EutE [8, 13]. Acetyl-CoA can be degraded to acetate with ATP production by the phosphotransacetylase EutD [14] and an alternative acetate kinase EutQ [15]. Alternatively, acetyl-CoA can be catabolized in the tricarboxylic acid cycle or the glyoxylate cycle, or used for lipid biosynthesis [8]. According to current models, EA is catabolized to acetate and ethanol in a molar ratio 1:1, via oxidative ATP-producing branch and reductive NAD+-regenerating branch [8]. Interestingly, previous studies showed that EA only confers a marked anaerobic growth advantage on *Salmonella enterica serovar Typhimurium* (*S. Typhimurium)* in the presence of tetrathionate, acting as an alternative electron acceptor via tetrathionate reductase [16]. And the mutant lacking the tetrathionate reductase showed a decreasing colonization capacity in a mouse colitis model, which points to a role for anaerobic electron transfer in EA catabolism contributing to growth of *S. Typhimurium* in the lumen of the inflamed intestine [16]. Anaerobic EA catabolism in *L. monocytogenes*, including possible roles for anaerobic respiration, has not been studied. Notably, *L. monocytogenes* lacks tetrathionate reductase, but anaerobic electron transfer with fumarate reduction via membrane-bound fumarate reductase [16] and recently described flavin-based extracellular electron transfer (EET) with Fe ^3+^ as an electron acceptor [2], could act as substitutes in BMC-dependent EA catabolism.

The enzymes of the indicated EA pathway are present in a bacterial microcompartment (BMC), and structural shell proteins that constitute the BMC building blocks are encoded by genes in the *eut* cluster [17]. BMCs consist of a capsule of semi-permeable shell proteins and encapsulated enzymes of metabolic pathways that liberate toxic intermediates in the lumen of the capsules [17-19]. In the formation of BMCs, so-called encapsulation peptides, 10–20 residue-long hydrophobic α-helices in the N-terminal of some core enzymes, play a key role in the encapsulation mechanism [20, 21]. In our previous study, evidence was provided for a role of BMC-dependent utilization of 1,2-propanediol in *L. monocytogenes* supporting anaerobic growth and metabolism, and encapsulated enzymes PduD PduL and PduP were found to contain encapsulation peptides [22]. Expression of the *eut* operon in *L. monocytogenes* is under the regulation of the two-component regulators EutVW sequestrated by a B12-binding riboswitch [23, 24]. Upregulation of the *eut* operon has been found in *L. monocytogenes* grown on vacuum-packed cold smoked salmon and in co-cultures with cheese rind bacteria, which suggests a possible role of *eut* operon in the adaptation of *L. monocytogenes* to available nutrient sources [25, 26]. The *L. monocytogenes eut* operon exhibited increased expression inside the host cell, and loss of one of the key enzymes, ethanolamine ammonia lyase (EutB), caused a defect in intracellular growth [27].

Here we show by using metabolic analysis, transmission electron microscopy and proteomics, that *L. monocytogenes* forms *eut* BMCs and utilizes EA as a carbon source in anaerobic conditions with end products acetate and ethanol in a molar ratio 2:1. We demonstrate that the resulting redox imbalance is compensated by the flavin-based EET system by comparative growth and metabolic analysis of *L. monocytogenes* wild type, a BMC mutant and a EET mutant. Our results suggest an important role of anaerobic BMC-dependent EA catabolism in the physiology of *L. monocytogenes*, with a crucial role for the flavin-based EET system in redox balancing.

## Results

### EA utilization facilitates anaerobic growth

To find out whether EA utilization stimulates anaerobic growth, we added EA (15 mM) to LB medium in line with previous works in *Salmonella* species [13, 28]. The culture medium also contained 20 nM vitamin B12 for activation of the *eut* operon in *L. monocytogenes* [23]. *L. monocytogenes* EGDe cultures grown in *eut*-induced condition (LB with EA and B12) reached significantly higher OD_600_ values after 8h of incubation compared to cultures in control conditions (LB and LB with B12) and in *eut* non-induced conditions (LB with EA) (Figure 1A). EA in *eut*-induced condition was fully utilized within the first 24h while no significant EA utilization was observed in *eut* non-induced conditions (Figure 1B). Notably, in *eut*-induced condition EA was converted into acetate and ethanol (Figure 1C and 1D), while in *eut* non-induced condition only acetate was produced conceivably originating from metabolism of other compounds in LB. From this comparative analysis on acetate production we derive a molar ratio acetate:ethanol of approximately 2:1. Taken together, the utilization of EA with the production of acetate and ethanol contributes to the anaerobic growth of *L. monocytogenes* EGDe in *eut* induced condition.

**Figure 1.**
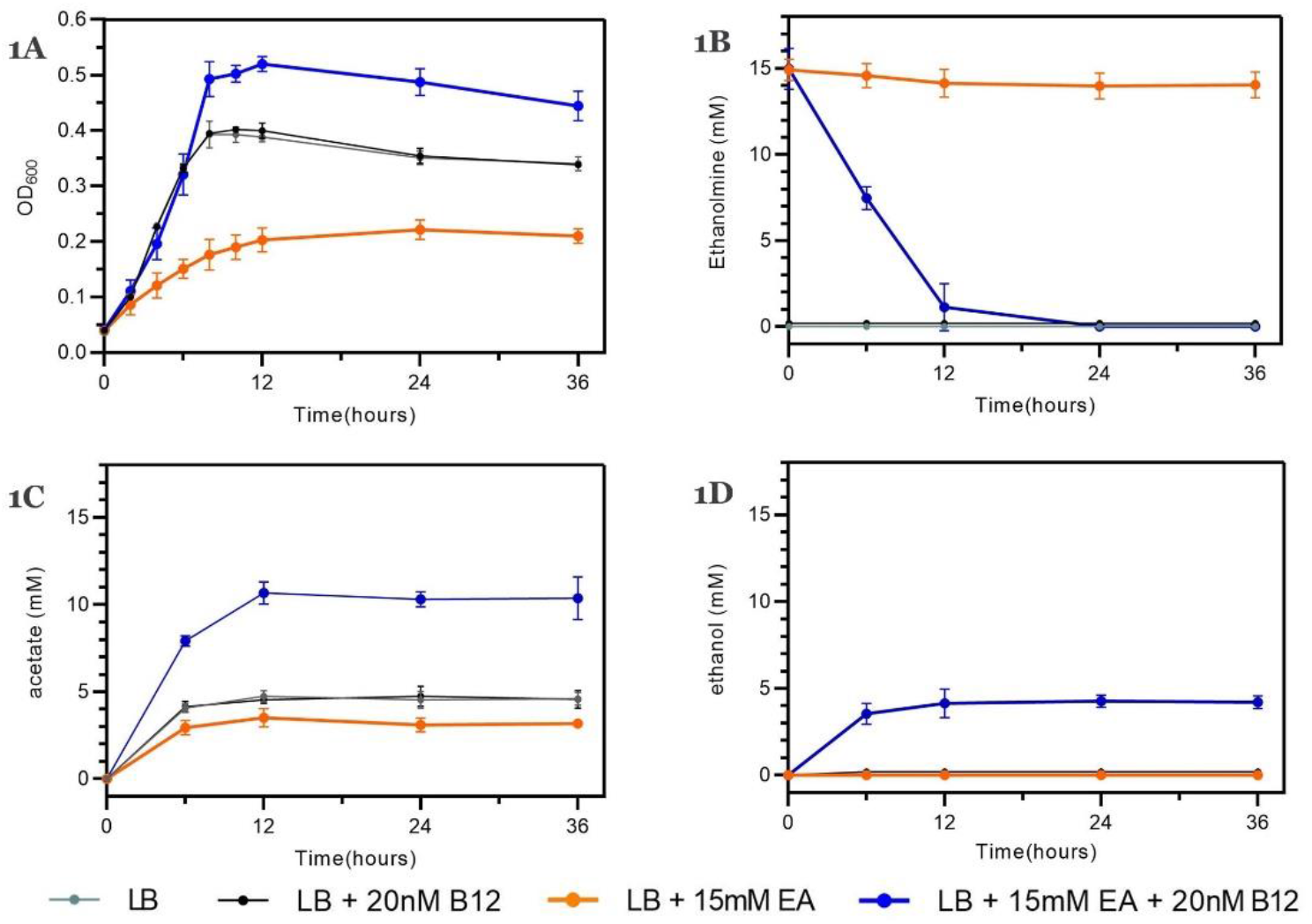
Anaerobic growth and EA catabolism of *L. monocytogenes* EGDe in LB medium. **(A)** Impact of EA and/or vitamin B12 on anaerobic growth of *L. monocytogenes* EGDe; **(B)** EA Utilization; **(C)** Acetate production; **(D)** Ethanol production. Lines represent different growth conditions; Results from three independent experiments are expressed and visualized as means and standard errors.

### Ratio of ethanol and acetate production

To clarify whether *L. monocytogenes* EGDe can utilize EA as a sole carbon source, we examined EA utilization and its impact on anaerobic growth in defined medium MWB in absent of any other carbon sources [29]. *L. monocytogenes* EGDe inoculated in *eut* induced condition (MWB with EA and B12) showed about 100-fold increase in cell counts after 24h while no significant increase in cell counts was observed in non-induced condition (MWB with EA) (Figure 2A). During the anaerobic growth of *L. monocytogenes* EGDe, 15 mM EA was converted into about 6.6 mM acetate and 3.4 mM ethanol in *eut* induced condition while no significant degradation of EA was observed in non-induced condition (Figure 2B, 2C and 2D). Calculation of the carbon mass balance, part of EA (approximately 5mM) is conceivably further catabolized in the glyoxylate cycle, or used for lipid biosynthesis via the intermediate acetyl-CoA [8]. The utilization of EA in *eut* induced condition in defined medium MWB provides evidence that EA can act as a sole carbon source supporting anaerobic growth of *L. monocytogenes* EGDe. Notably, the observed molar ratio acetate:ethanol of 2:1 suggests an apparent redox imbalance due to higher flux via the NADH and ATP-producing acetate branch, and reduced regeneration of NAD+ in the ethanol branch resulting in a surplus of NADH.

**Figure 2.**
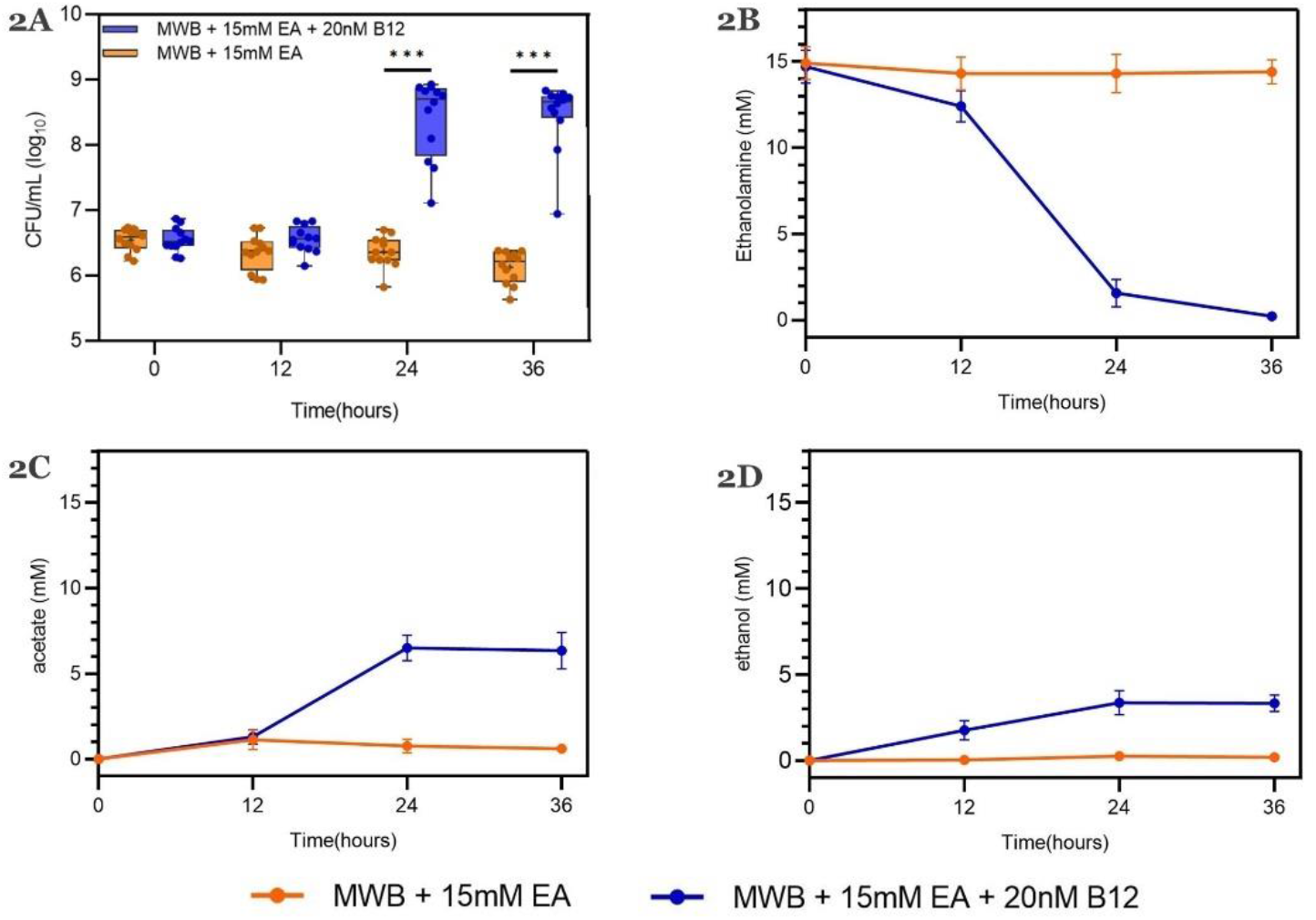
Anaerobic growth and EA catabolism of *L. monocytogenes* EGDe in MWB defined medium with EA as sole carbon source. **(A)** Impact of EA and/or vitamin B12 on CFUs of *L. monocytogenes* EGDe; Results from three independent experiments with four technical repeats are expressed as mean and standard errors. Statistical significance is indicated (***, P<0.001; Holm-Sidak T-test) **(B)** EA Utilization; **(C)** Acetate production; **(D)** Ethanol production. Lines represent different growth conditions; Error bars in (B, C, D) indicate three independent experiments expressed as mean and standard errors.

### Upregulated expression of *eut* operon including enzymes and structural shell proteins

In order to study the expression of the *eut* operon and the way it is imbedded in cell physiology, we performed proteomics to compare *L. monocytogenes* cells grown in LB and MWB medium under *eut* induced conditions and non-induced conditions. Analyses of the complete list of identified proteins, proteins’ expression levels and subsequent t-test results and p-values are shown in (<u>Supplementary Table 2</u>) for LB and (<u>Supplementary Table 3</u>) for MWB grown cells. For LB, we identified 1891 total proteins where 161 are upregulated more than two-fold and 229 proteins downregulated more than two-fold in *eut* induced condition compared to non-induced condition (Figure 3A). Among these 161 upregulated proteins, the top 15 proteins are all encoded in the *eut* operon, i.e., Eut GABCLKEMTDNHQ, lmo1183 and 1185 while the other two proteins Eut VW [30] involved in *eut* operon regulation are also included. For MWB, 1736 proteins were identified of which 253 proteins are upregulated more than two times and 162 proteins downregulated more than two times in *eut* induced condition compared to non-induced condition (Figure 3B). In line with LB data, the top 15 proteins are all encoded in the *eut* operon. Analysis of the upregulated proteins in LB and MWB shows 50 proteins that overlap between the conditions pointing to a prominent role in *eut* induced conditions compared to noninduced conditions. Among these 50 proteins, 17 proteins are linked to *eut* of which 16 proteins are encoded in the *eut* operon and one gene *dra* (deoxyribose-phosphate aldolase) is predicted to have an interaction with the acetaldehyde dehydrogenase *eutE* (lmo1179) (according to STRING, Figure 3C, <u>Supplementary Table 4</u>). Another group of overlapping genes are potential ABC transporters, including *lmo2751-Imo2752* (ABC transporter ATP-binding protein) [31], and *ilvC* (*lmo1986*, NADP+ based Ketol-acid reductoisomerase)[32]. The analysis of downregulated proteins in LB and MWB showed an overlap of 48 proteins with 18 proteins linked to vitamin B12 biosynthesis which indicates that adding vitamin B12 in the medium represses the genes involved in vitamin B12 biosynthesis (Figure 3D, Supplementary Table 5). Overlap in downregulated proteins also showed an enrichment in phospholipase (*plcA* and *plcB*), cell wall relevant (*lmo2691* and *lmo0695*), cell division relevant (*dilvB, rpmG1* and *lmo0111*) and zinc containing dehydrogenase (*lmo2663, lmo2664* and *lmo2097*). To summarize, the significant upregulation of the *eut* operon at proteomic level including structural shell proteins strongly supports that BMC-dependent EA utilization is processed by enzymes and structural shell proteins of the *eut* operon.

**Figure 3.**
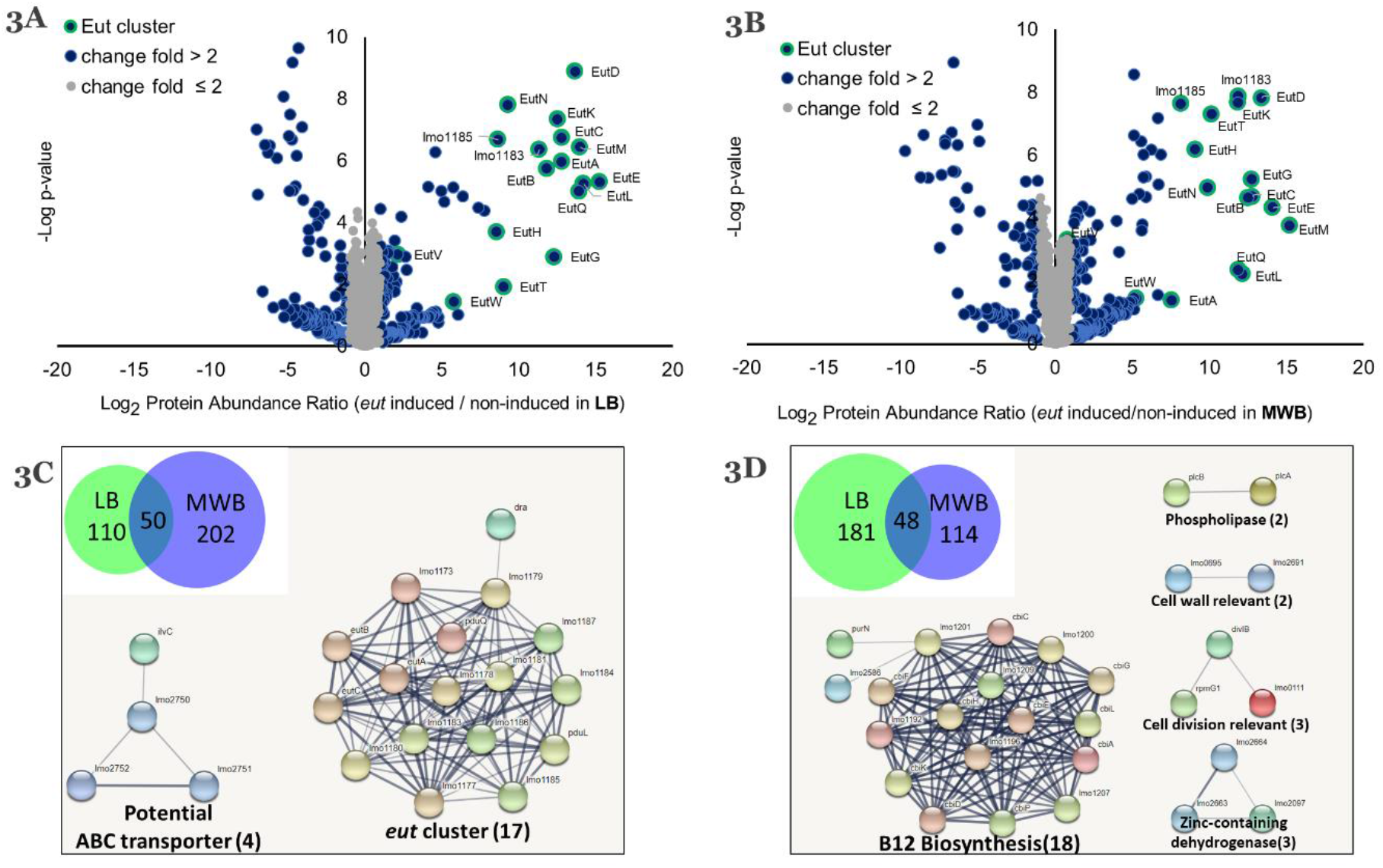
Proteomics analysis of *eut* induced and non-induced *L. monocytogenes*EGDe in LB medium and MWB defined medium. Proteomic volcano plot of *eut* induced cells (EA + B12 added) compared to noninduced cells (EA added only) in LB medium **(A)** and MWB medium **(B)**. Venn diagram of 50 overlapping upregulated proteins **(C)**, and Venn diagram of 48 overlapping downregulated proteins **(D)**, for LB medium (green) and MWB medium (blue) and corresponding STRING protein-protein interactions. Nodes represent proteins and lines represent interactions.

### BMC structures support BMC-dependent EA catabolism

To further confirm the presence of BMCs in *eut* induced cells, we used transmission electron microscopy (TEM) and compared thin sections of both eut-induced and noninduced *L. monocytogenes* EGDe cells. The *eut* induced cells clearly contain BMC-like structures with an approximate diameter of 50–80 nm, which are not present in non-induced cells (Figure 4B). Notably, the identified structures strongly resemble TEM pictures of BMCs in *S. Typhimurium and E. coli* [33, 34], and that of recently reported BMCs found in *pdu*-induced *L. monocytogenes* [22]. Taken the metabolic, proteomic, and TEM data together, we conclude that the cytosolic BMC-like structures in *L. monocytogenes* EGDe are involved in EA utilization under anaerobic conditions. Hereby, based on the knowledge of the *eut* operon, we propose a model of the BMC-dependent EA catabolism. The *eut* operon in *L. monocytogenes* EGDe contains 17 genes and is most likely under the regulation of two-component regulators EutVW sequestrated by binding of vitamin B12 to the riboswitch rli55 [23]. In front of EutVW, we found a previously predicted *cre* site for the binding of carbon control protein CcpA pointing to catabolite repression control of the *L. monocytogenes eut* cluster [35]. Five *eut* genes, *eut LKMN* and *lmo1185*, are predicted to be structural shell proteins of the BMC (Figure 4A). EutK and EutM are the hexameric shell protein (BMC-H) consisting of one Pfam00936 domain while EutL and lmo1185 are the trimeric shell proteins (BMC-T) with two fused Pfam00936 domains (<u>Supplementary Figure 1</u>). Notably, the trimeric assembly of lmo1185 forms a flat approximately hexagonally shaped disc with a central pore that is suitable for a [4Fe-4S] cluster [1]. Furthermore EutN is the pentameric shell protein (BMC-P) consisting of one Pfam03319 domain(<u>Supplementary Figure 1</u>). BMC is assembled by these three types of shell proteins: BMC-H, BMC-T and BMC-P [17, 18]. Based on the previous BMC-dependent EA catabolism model in *S. Typhimurium* [7], we propose a model for *L. monocytogenes* with putative encapsulation peptides supporting recruitment of selected eut enzymes to the BMC (Figure 4C). We predict a specific hydrophobic α-helix in the N terminus of the encapsulated enzyme EutC, EutD and EutE (<u>Supplementary Figure 2</u>), similar to that of the previously described BMC encapsulated proteins in *S. Typhimurium* and for *pdu* BMCs in *L. monocytogenes* [20, 22]. EA is split into acetaldehyde and ammonia by ethanolamine ammonia lyase EutBC[8, 11, 12]. Acetaldehyde can be converted into ethanol by the alcohol dehydrogenase EutG[10], or into acetyl-CoA by the acetaldehyde dehydrogenase EutE[8, 13]. Acetyl-CoA can be converted into acetyl-phosphate and subsequently acetate and ATP production by the phosphotransacetylase EutD[14] and an alternative acetate kinase respectively EutQ[15].

**Figure 4.**
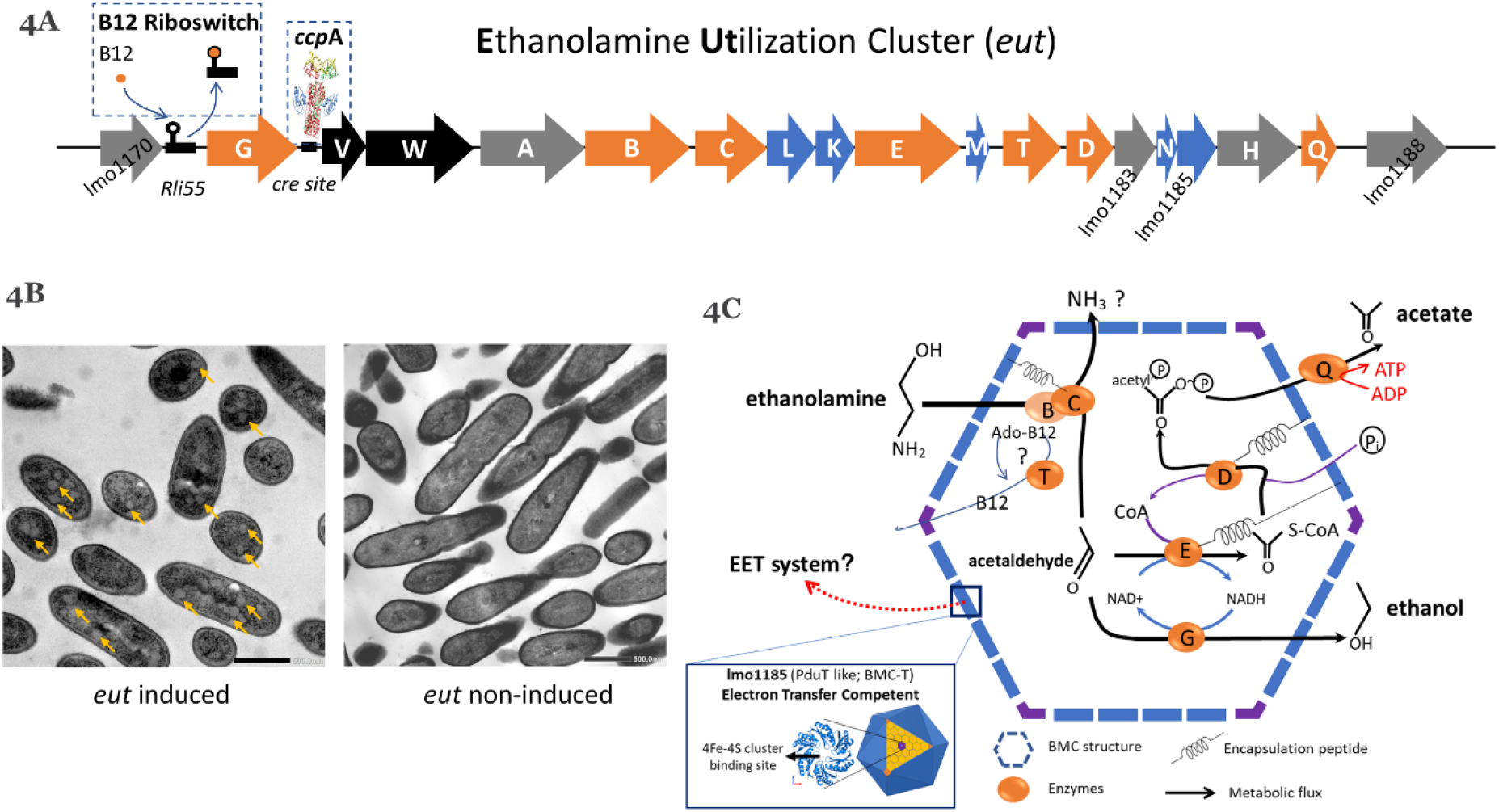
Overview of BMC-dependent EA catabolism model of *L. monocytogenes*. **(A)** Analysis of the *eut* operon (<u>Details in Supplementary Table 1</u>). Characters in orange represent Eut enzymes, in blue represent BMC shell proteins, in black represent the two component regulation system, and in grey represent unannotated proteins; B12 riboswitch and CcpA *cre* binding site are indicated. **(B)** TEM visualization of BMCs in *eut* induced (left; yellow arrows point to BMCs) and noninduced cells (right). **(C)** Model of BMC-dependent EA catabolism. EutBC ethanolamine ammonia lyase, EutD phosphotransacetylase, EutE acetaldehyde dehydrogenase, EutG alcohol dehydrogenase, EutQ acetate kinase, EutT corrinoid cobalamine adenosyltransferase, with putative encapsulation peptides indicated. Zoom-in part shows the prediction of potential [4Fe-4S] cluster binding site in Eut shell protein lmo1185, that is highly similar to that previously reported for PduT shell protein by Pang et al. [1]. See text for details.

### Flavin-based EET linked with BMCs maintains redox balance of EA catabolism

The observed unbalanced production of acetate and ethanol in a 2:1 molar ratio suggests a surplus of NADH, and this requires additional NAD+ regeneration reactions to restore the redox balance. As discussed in previous studies, BMC is a redox-replete compartment which can generate reductants internally or facilitate the transfer of electrons from the cytosol across the shell [8, 36]. Recently it has been shown that *L. monocytogenes* uses a distinctive anaerobic flavin-based EET mechanism to deliver electrons to iron (Fe^3+^) or to fumarate via membrane-bound fumarate reductase [2]. In our study, MWB defined medium indeed contains ferric citrate and flavin, and proteomic analysis of *L. monocytogenes* EGDe grown in this medium identified protein lmo2637 encoding a EET-linked lipoprotein PplA and protein lmo2638 encoding a EET-linked NADH dehydrogenase Ndh2 [2] (<u>Supplementary Table 3</u>). Next, we tested the hypothesis that anaerobic EET could play a role in BMC-dependent EA utilization using *L. monocytogenes* 10403S wild type and mutants. Notably, *eut* induced *L. monocytogenes* 10403S showed a similar growth benefit from utilizing EA as *eut* induced *L. monocytogenes* EGDe (Figure 5A and Figure 5B). Moreover, the mutant strains *L. monocytogenes* 10403S Δ*eut*B, lacking ethanolamine ammonia lyase, and Δ*ndh*2 lacking EET-linked NADH dehydrogenase are both impaired for EA utilization (Figure 5B), suggesting an involvement of the EET system in EA utilization and thereby a link with BMCs. Next, we used a ferrozine-based colorimetric assay the determine electron transfer between BMC and EET following *L. monocytogenes* EA utilization in MWB medium. In this assay, electrons generated from EA catabolism are transferred to Fe^3+^ generating Fe^2+^, where binding of ferrozine to the differently charged Fe molecules results in a colorimetric change from yellow brown to fuchsia, respectively. The colorimetric changes were shown for *L. monocytogenes* EGDe and *L. monocytogenes* 10403S in *eut* induced condition following EA utilization while *eut* mutant strain *L. monocytogenes* 10403S Δ *eut*B and EET mutant strain *L. monocytogenes* 10403S Δ *ndh*2 showed no colorimetric changes (Figure 5C). OD_562_ measurements indicated a significantly higher ferric iron reductase activity in *eut* induced conditions compared to non-induced conditions of *L. monocytogenes* EGDe and *L. monocytogenes* 10403S (Figure 5D). Taken together, these results provide evidence for a link between eut BMCs and the EET system via regeneration of NAD+ by NADH oxidation in the EET using Fe^3+^ as an electron acceptor (Figure 5E). The action of the EET as an alternative reductive pathway next to the ethanol branch, thus stimulates anaerobic growth by enhanced flux via the ATP generating acetate branch in BMC-facilitated EA catabolism.

**Figure 5.**
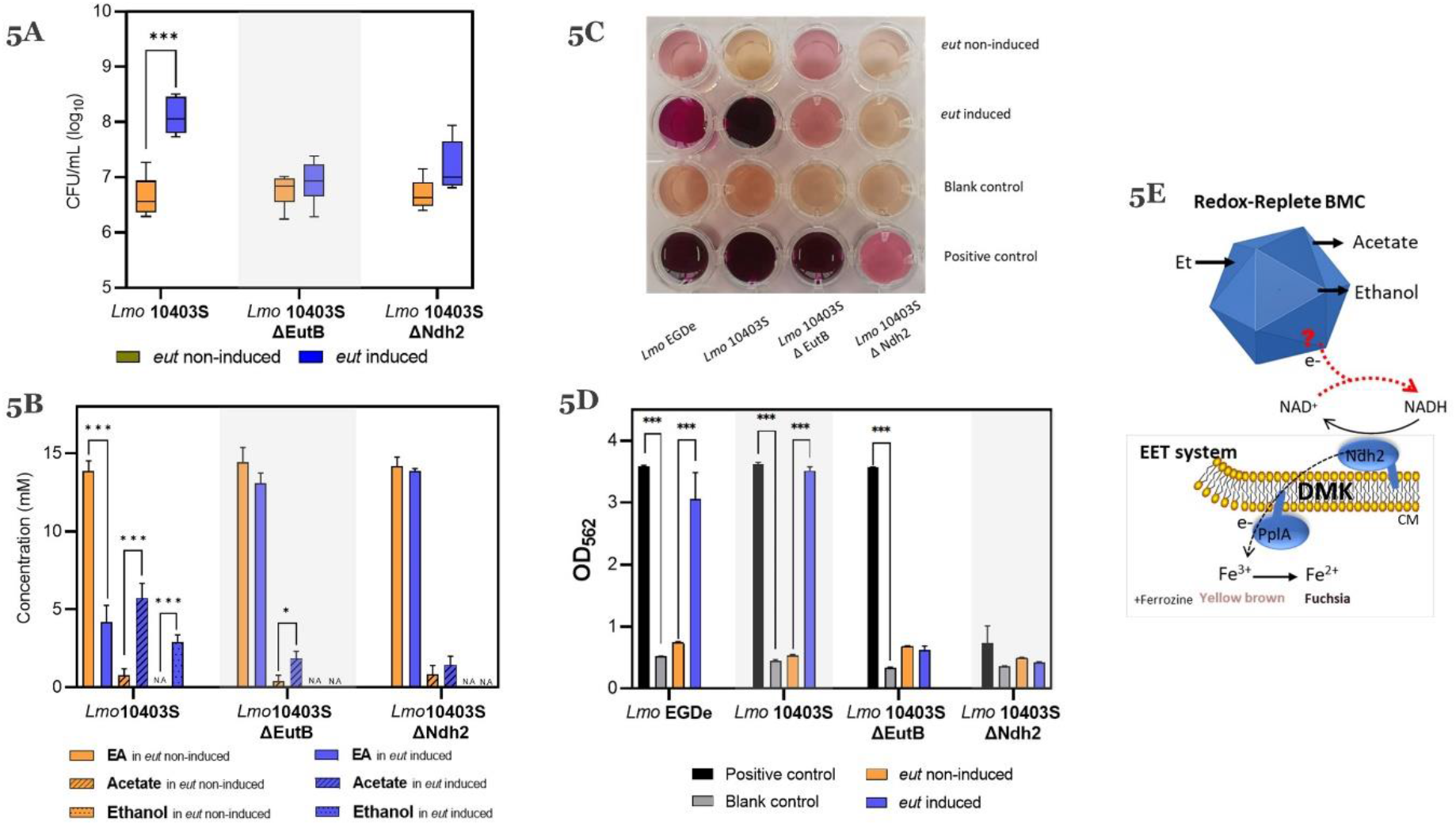
BMC-dependent EA catabolism couples to flavin-based EET. **(A)** Impact of EA and/or vitamin B12 on CFUs. **(B)** EA catabolism. Experiments in (A) and (B) were performed with *L. monocytogenes* 10403S and mutant strains grown anaerobically for 24h with initial 6.5 ± 0.1 log10 CFU/mL inoculation in MWB, with 15mM EA and 20nM B12 (*eut*-induced), or with 15mM EA (non-induced). **(C)** Colorimetric change of ferric reductase assay. **(D)** OD_562_ measurements of ferric reductase assay. Experiments in (C) and (D) were performed with *L. monocytogenes* EGDe and 10403S and mutant strains grown anaerobically for 24h in MWB, with 15mM EA and 20nM B12 (*eut*-induced), or with 15mM EA (non-induced), 15 mM glucose (positive control) and with no added substrate (blank control). Results in (A) (B) and (D) from three independent experiments are expressed as mean and standard errors. Statistical significance is indicated (***, P<0.001; *, P<0.05; Holm-Sidak T-test). (E) Proposed model of the electron transfer from BMC to EET. Blue geometric block represents BMC. CM represents cytoplasmic membrane; Ndh2, PplA and DMK represent EET with Fe^3+^ as electron acceptor (See text for details; figure 5E adapted from [2]).

## Discussion

This study provides evidence for the activation of BMC-dependent ethanolamine (EA) utilization in *L. monocytogenes* in anaerobic conditions in LB and MWB medium containing EA and vitamin B12. By using metabolic analysis, proteomics and electron microscopy we demonstrated the formation of BMCs in conjunction to EA catabolism with the production of acetate and ethanol in a 2:1 molar ratio. Selected genes in the *eut* operon encode structural shell proteins that form the respective *eut* BMC [17]. Previous studies showed that BMCs for EA catabolism in *S. Typhimurium* and *E. faecalis*, are composed of five structural shell proteins, EutS, EutM, EutK, EutL, and EutN [8, 17, 19]. Notably, *L. monocytogenes eut* operon also encodes five putative shell proteins, EutM, EutK, EutL, EutN and lmo1185, and combined with visualization of BMC structures by TEM and our proteomic data, we conclude that *eut* BMCs are composed of these five indicated structural shell proteins. Apparently, EutS is not essential for BMC assembly in *L. monocytogenes*. EutS is hexameric BMC shell protein with a Pfam00936 domain, and it is conceivable that function of EutS is taken over by EutK and/or EutM in *L. monocytogenes*, since both are also hexameric BMC shell proteins with a Pfam00936 domain (<u>Supplementary Figure 1</u>).

EA is a valuable carbon source for *L. monocytogenes* to outcompete other bacteria unable to utilize EA in food environments [25, 26] or human gastrointestinal tract where EA is abundant [7, 8]. Our results in defined medium reveal that *L. monocytogenes* can utilize EA as sole carbon source via a BMC-dependent *eut* pathway (Figure 2). Encasing the pathway inside BMCs is essential, since this prevents the toxic acetaldehyde intermediate to damage proteins and RNA/DNA in the cytoplasm [17, 37]. The generated reductants are oxidized inside the BMC, while it has been hypothesized that electrons may also be shuttled to the cytosol via specific shell proteins acting as redox carriers [8, 17, 18, 36]. Our metabolite analysis showed enhanced flux via the ATP generating acetate branch in *L. monocytogenes*, and a reduced flux via the NAD+ regenerating ethanol branch in a 2:1 molar ratio resulting in surplus NADH. In support of the previously mentioned putative electron shuffling from BMCs to the cytosol, using wild type, EutB and Ndh2 mutants, we identified a link between *L. monocytogenes eut* BMCs activity and the recently discovered flavin-based EET system [3]. Suggested electron acceptors, fumarate and iron, are conceivably present in the human intestine and in host cells and have been reported to contribute to *L. monocytogenes* virulence [2, 16, 38, 39]. The identified *L. monocytogenes eut* Lmo1185 shell protein belongs to the class of trimeric shell proteins (BMC-T) with two fused Pfam00936. We hypothesize that this predicted PduT-like shell protein with a [4Fe-4S] cluster [1], acts as a redox carrier in this process. Further studies are required to elucidate the role of this *eut* BMC shell protein as redox carrier. Taken together, our results provide evidence of anaerobic EA catabolism in *L. monocytogenes* driven by BMC expression and function with a crucial role for the flavin-based EET system in redox balancing.

## Materials and Methods

### Strains, Culture Conditions, and Growth Measurements

All *L. monocytogenes* strains used in this study were shown in (<u>Supplementary Table 6</u>). *L. monocytogenes* strains were anaerobically grown at 30°C in Luria Broth (LB) medium and defined medium MWB[29]. LB and MWB were supplemented with 15mM EA and/or 20nM vitamin B12 [23]. Anaerobic conditions were achieved by Anoxomat Anaerobic Culture System with the environment 10% CO2, 5% H2, 85% N2. LB and MWB with 15 mM EA and 20 nM vitamin B12 were defined as *eut*-induced condition, while LB and MWB with 15 mM EA were defined as non-induced condition. OD_600_ measurements in LB were performed every 2 h during the first 12 h of incubation and at 24 and 36 h. Plate counting in MWB to quantity Colony Forming Units (CFUs) was performed every 12 h from 0h to 36h.

### Construction of strain *L. monocytogenes* 10403S Δ*eutB*

*L. monocytogenes* Δ*eut*B strain was derived from wild-type 10403S (DP-L6253). Gene deletions were generated by allelic exchange using the plasmid pKSV7 [40] with chloramphenicol resistance gene. Primers used for Δ*eut*B fragment construction and validation of deletion are given in (<u>Supplementary Table 7</u>). Fragment A and fragment B flanking *eutB* gene were ligated into plasmid using Gibson assembly and cloned in Top10 *E. coli*. Plasmid construct was verified by Sanger sequencing. Plasmid was then transformed into SM10 *E. Coli*. Transconjugation was performed to integrate plasmid using restrictive temperature of 42°C and colonies that were resistant to streptomycin (200 μg/mL) and chloramphenicol (7.5 μg/mL) were selected. This was followed by passaging in Brain-Heart Infusion at 37°C with shaking, diluting 1:1000 every 8 h. Mutant was obtained by screening for chloramphenicol sensitive colonies followed by colony PCR using validation primers.

### Analysis of metabolites for EA catabolism

After centrifugation, the supernatants of the cultures were collected and filtered with 0.45 μm syringe filter for the measurements. EA(mono-ethanolamine) with 1:1 dilution in ethanol was measured by Gas Chromatography with Flame Ionization Detection (GC-FID) while ethanol and acetate were directly measured by High Pressure Liquid Chromatography (HPLC). The experiment was performed twice with three technical replicates per experiment. Additionally, the standard curves of EA, ethanol and acetate were measured in the concentration range of 0.1, 1, 5, 10, 15mM. HPLC was performed as described before[22]. GC-FID conditions for the final method were as follows: 0.3-μL injection, 260 °C injector temperature, 1177 injector, SGE focus liner, 1:10 split, 335 °C detector temperature, and electronic flow controller delivering 2.8 mL/min helium carrier gas with a 2.0 psi pressure pulse for 0.25 min after injection. The retention time of EA was at 11 min, and the total run time was 50 min [41].

### Proteomics

*L. monocytogenes* EGDe cultures were anaerobically grown at 30°C in *eut* induced and in non-induced conditions. Samples were collected at 12 h for LB and 24 h for MWB. Then samples were washed twice with 100 mM Tris (pH 8). About 10 mg wet weight cells in 100 μl 100 mM Tris was sonicated for 30 s twice to lyse the cells. Samples were prepared according to the filter assisted sample preparation protocol (FASP) [42]. Each prepared peptide sample was analyzed by injecting (18 μl) into a nanoLC-MS/MS (Thermo nLC1000 connected to a LTQ-Orbitrap XL) [22]. LCMS data with all MS/MS spectra were analyzed with the MaxQuant quantitative proteomics software package as described before [43]. A protein database with the protein sequences of L. monocytogenes EGDe (ID: UP000000817) was downloaded from UniProt. Filtering and further bioinformatics and statistical analysis of the MaxQuant ProteinGroups file were performed with Perseus [44]. Reverse hits and contaminants were filtered out. Protein groups were filtered to contain minimally two peptides for protein identification of which at least one is unique and at least one is unmodified. Also, each group (eut-induced and non-induced control) required three valid values in at least one of the two experimental groups. The volcano plot was prepared based on the Student’s t-test difference of *eut*-induced/non-induced.

### Transmission Electron Microscopy

*L. monocytogenes* EGDe cultures were grown anaerobically at 30°C in *eut*-induced or non-induced conditions. Samples were collected at 12 h of incubation for LB (early stationary phase). About 10 μg dry cells were fixed for 2 h in 2.5% (v/v) glutaraldehyde in 0.1 M sodium cacodylate buffer (pH 7.2). After rinsing in the same buffer, a post-fixation was done in 1% (w/v) OsO4 for 1 h at room temperature. The samples were dehydrated by ethanol and were then embedded in resin (Spurr HM20) 8 h at 70°C. Thin sections (<100 nm) of polymerized resin samples were obtained with microtomes. After staining with 2% (w/v) aqueous uranyl acetate, the samples were analyzed with a Jeol 1400 plus TEM with 120 kV setting.

### Ferrozine assay of ferric iron reductase activity

*L. monocytogenes* cells grown overnight in LB medium were washed with PBS twice, normalized to an OD_600_ of 0.2, and resuspended in fresh MWB medium supplemented with 50 mM ferric ammonium citrate for anaerobic inoculation. *L. monocytogenes* Wild Type and mutant strains grew anaerobically for 24h in MWB, with 15mM EA and 20nM B12 (*eut*-induced), or with 15mM EA (non-induced), 15 mM glucose (positive control) and with no added substrate (blank control). Assays were initiated by adding 100 μL of MWB cultures from 24 h anaerobic inoculation to 100 μL demi water with 4 mM ferrozine and then spectrophotometrically measured by OD_562_ as described before [2]. OD_562_ measurements were made immediately after the initial mixture of MWB cultures and Ferrozine.

### Bioinformatics Analysis

Secondary Structure of N Terminal Peptides. The N terminal secondary structures of all *eut* genes were determined by a neural network secondary structure prediction called Jpred4 [45] as described before [22]. The input to the Jpred 4 online server1 was the 50 N-terminal amino acids of each protein. Jnetconf: confidence estimation for the prediction with high scores indicating high confidence. Jnetsol25: solvent accessibility, where B means buried and ‘-’means non-buried at 25% cut-off.

Venn analysis and STRING networks analysis of proteins. The protein IDs of significantly changed proteins from <u>Supplementary Table 4 and 5</u> were uploaded to the BioVenn online server[46] taking the default setting to generate Venn diagrams. Overlapping proteins from the Venn diagram were transferred to the STRING online server[47] for multiple proteins analysis of functional interaction using sources such as co-expression, genomic neighborhood and gene fusion.

## Acknowledgments

We thank the Wageningen Electron Microscopy Centre for TEM support. We thank Kees van Kekem in Wageningen Food & Biobased Research for GC-FID support. ZZ was supported by a grant from the China Scholarship Council.

## Author Contributions

ZZ, RN, and TA designed the experiments. ZZ performed experiments. SB and ZZ performed proteomics measurements. VB constructed the EutB mutant strain. SL provided input with respect to the BMC-EET system and supported EET experiments. RN and TA supervised the research. ZZ, RN, and TA analyzed data and wrote the manuscript with improvements from SB, SL and ES.

